# Targeted RNA degradation of HIV-1 TAR element at 5’UTR by CRISPR-Cas13X blocks HIV-1 lytic reactivation

**DOI:** 10.64898/2026.10.14.759209

**Authors:** Thurbu T. Lepcha, Dawei Zhou, Guillaume N. Fiches, Nithya Paruchuri, Jinshan He, Kateepe A.S.N. Shanaka, Yan Liu, Suha Eleya, Netty G. Santoso, Jian Zhu

## Abstract

Human immunodeficiency virus type 1 (HIV-1) latency remains a major barrier to a cure. Here, we developed a hypercompact CRISPR-Cas13X.1 system to target the HIV-1 5′ untranslated region (5′UTR), focusing on the transactivation response (TAR) RNA element required for Tat-dependent transcription. Following validation of a single-vector system, we screened CRISPR RNAs (crRNAs) targeting distinct 5′UTR elements and identified crRNA-TAR-2 as the most effective inhibitor of Tat-dependent reporter activity. Mechanistic analyses demonstrated that TAR targeting disrupted Tat–TAR interactions. In HIV-1 infection models, Cas13X.1-mediated TAR targeting reduced HIV-1 Gag precursor (p55) and capsid (p24) protein expression, as well as gag and tat transcript levels, accompanied by activation of the PKR–eIF2α antiviral pathway. TAR targeting also suppressed viral gene expression and reactivation in multiple latency models and reduced viral load in HIV-1-infected primary peripheral blood mononuclear cells (PBMCs). These findings establish HIV-1 TAR RNA as a target for Cas13X.1-mediated antiviral activity and support RNA-targeting CRISPR systems as a potential strategy to suppress HIV-1 infection and reactivation.

## INTRODUCTION

Human immunodeficiency virus type 1 (HIV-1), a member of the *Retroviridae*, carries a single-stranded RNA genome that is reverse transcribed into double-stranded DNA and integrated into the host-cell genome to establish a provirus and global burden to more than 34 million people (1–3). Although combination antiretroviral therapy (cART) effectively suppresses HIV-1 replication, it cannot eradicate infection because of persistent latent viral reservoirs, while the error-prone nature of HIV-1 reverse transcriptase contributes to viral genetic diversity and the potential emergence of drug-resistant variants (4–6).The RNA structures from the HIV-1 genome provide valuable insights into the mechanisms underlying the viral replication cycle. Moreover, these structures serve as models for designing novel therapeutic approaches (7).

Strategies to eliminate or silence HIV-1 have generally pursued one of two levels of intervention, permanent modification of the integrated proviral DNA or targeted degradation of viral RNA. RNA interference (RNAi) and CRISPR-Cas9 technologies are two widely used approaches for reducing gene expression (8). However, RNAi-based gene silencing has been associated with significant off-target effects, which may limit its specificity and therapeutic applications (9). Although CRISPR-Cas9-based genome editing has been widely explored for excising or disrupting the HIV-1 provirus, but this DNA-targeting approach carries an inherent risk of off-target genomic modification and requires rigorous safety validation before clinical translation (10,11).Moreover, HIV-1 proviral quasispecies are genetically heterogeneous, limiting uniform access by editing machinery, unedited copies continue to transcribe virus, and NHEJ-mediated repair of Cas9-induced double-strand breaks can generate indel-associated escape variants (12,13). These limitations have motivated RNA-level interventions, which suppress viral gene expression without permanently altering the genome and act independently of integration site.

Cas13 was first identified through the characterization of C2c2 (now designated Cas13a), an RNA-guided ribonuclease from *Leptotrichia shahii*. C2c2 uses a single crRNA to recognize and cleave complementary single-stranded RNA targets, providing bacterial defense against RNA phage, and its programmability for targeted mRNA knockdown established the foundation for RNA-targeting CRISPR applications (14). Cas13 effectors belong to the Type VI CRISPR-Cas family, which is unique among characterized CRISPR-Cas systems in possessing RNA-guided RNase activity without any accompanying DNase activity, distinguishing it from DNA-targeting effectors such as Cas9 and Cas12 (15–17). Type VI systems target single-stranded RNA (ssRNA) and comprise six identified subtypes (VI-A–D, VI-X, and VI-Y), five of which have been functionally characterized with their respective signature effectors: Cas13a, Cas13b, Cas13d, Cas13X, and Cas13Y (15–17). Among these, Cas13d, commonly referred to as CasRx, attracted particular attention for its compact size of 930 amino acids, making it the first Cas13 ortholog amenable to single-vector adeno-associated virus (AAV) delivery (18). Despite this early promise, CasRx has since been associated with significant collateral RNA cleavage, including a report of lethal 28S rRNA degradation in mice (19) and evidence of collateral activity in human cells (20) , with its overall specificity profile remaining a subject of ongoing debate (21).

These concerns motivated our selection of Cas13X.1 (also known as Cas13bt3), a smaller and reportedly more specific Cas13 ortholog, for targeting the HIV-1 TAR element in this study (16). Cas13X.1 is a hypercompact effector capable of programmable RNA editing and knockdown in mammalian cells, combining high interference efficiency with low collateral activity and offering a more favorable specificity profile than CasRx for RNA-targeting applications (16,22)

The 5′-untranslated region (5′-UTR) of the HIV-1 genomic RNA is organized into several distinct structural domains, including the transactivation response element (TAR; nt 1–58), the polyadenylation signal hairpin (polyA; nt 59–105), the primer-binding site (PBS; nt 126–224), and the packaging signal (Psi; nt 229–335)(23) . As the most conserved region of the HIV-1 genome, this highly structured 5′-UTR participates in multiple stages of the viral replication cycle (24). Among its constituent elements, the TAR stem-loop is the best characterized, functioning as the binding platform through which the viral Tat protein activates transcription. Present at the 5′ terminus of HIV-1 transcript, TAR adopts a conserved RNA hairpin structure containing a trinucleotide pyrimidine bulge and an apical loop; together, these features confer high-affinity Tat binding and enable recruitment of the host P-TEFb elongation complex, a step indispensable for viral transactivation (25–30) . Despite selective pressure favoring its disruption, TAR’s structure remains remarkably conserved across primate lentiviruses, underscoring its functional constraints and establishing it as a long-standing target for therapeutic strategies aimed at interrupting the Tat-TAR-P-TEFb axis (29,31) . Given that TAR forms at the very start of nascent viral transcripts, RNA-guided Cas13 effectors particularly PFS-flexible orthologs like Cas13X.1could in principle be programmed to cleave this region co-transcriptionally, offering a modern nuclease-based extension of classical TAR-targeting approaches.

Although Cas13-based HIV-1 inhibition has been demonstrated using Cas13a and RfxCas13d, targeting coding regions such as *gag*, *pol*, and cPPT (32,33), a strategy specifically targeting the HIV-1 5′UTR remains unexplored. Moreover, no study to date has applied Cas13X.1, at 775 amino acids, the smallest known Cas13 ortholog and well suited for single-vector AAV delivery (16,18), to HIV-1 RNA targeting. Here, we designed CRISPR RNAs (crRNAs) against structural elements within the HIV-1 5′UTR, focusing on the TAR RNA element, and evaluated knockdown efficiency, disruption of the Tat-TAR interaction, inhibition of HIV-1 replication, and induction of antiviral activity through the PKR-eIF2α pathway.

## MATERIAL AND METHODS

### Reagents

Following antibodies are used, eIF2α (D-3) sc-133132, p-eIF2α (S51) CST-9721S, PKR CST#3072S, p-PKR (Invitrogen) Ref# 44-668G (Thr 451). Anti-CD3 (Cat. #16003785) and anti-CD28 (Cat. #16029805) antibodies were purchased from Ebioscience. Recombinant human TNF-α (Cat. # 554618) was purchased from BD Biosciences. Transfection reagents Fugen 6 and Turbofect was from Promega and Invitrogen respectively. Susfexin (Lamda Biosciences, cat no. ) was used for the transfections of suspension cells according to the manufacture protocol. HIV-1 IIIB and HIV-1 NL4-3 strains were required from the NIH AIDS reagent program.HIV-1 Gag p24 IgG1 monoclonal antibody was produced from the hybridoma cell line (NIH AIDS reagent program). The pQCXIP-HIV-Tat was constructed by subcloning C terminal Flag tag fused HIV-1 Tat into the pQCXIP retroviral empty vector (Clontech) using NotI and BamHI sites (34)

### Cas13X.1 crRNA design and plasmid construction

We designed crRNAs targeting the HIV-1 5’UTR, using the web tool CHOPCHOP. Pre-insertion sequences for the corresponding gRNAs were synthesised by IDT and cloned into a vector using the Zhang Lab protocol (35). We also designed and cloned the mCherry targeting crRNA in Cas13X.1 backbone plasmid to confirm the Cas13X.1 activity. The Cas13X.1 backbone plasmid (171379, Addgene) was digested with BbsI-HF enzyme in NEBuffer at 37°C for 4 hours. The larger fragment after digestion was extracted using the Gel extraction kit () . Ligation was performed with T4 DNA ligase () at 16°C overnight, followed by transformation of the ligated mixture into chemically competent DH5α cells. Single colonies were selected on ampicillin-resistant LB agar plates. Finally, sequence analysis was confirmed by Sanger sequencing.

### Cloning of HIV-1 5’UTR and Luciferase assay

The 5’-UTR of HIV-1 (356 bp) was cloned in the reporter vector pSicheck-2 (Addgene), harbouring the dual firefly and Renilla luciferase gene. The 5′UTR of HIV-1 was amplified from the pNL4-3 plasmid, using the sense and antisense primers 5’-CGG GCTAGC GCCTGTACTGGGTCTCTCTG-3’(A), and 5’-CCG GCTAGC CGCTTAATACCGACGCTCTC-3’(B)′.The resulting PCR product size was confirmed in the agarose gel, and cloned in the NheI sites of pSicheck-2 and sequenced using sanger sequencing. Cas13X.1 overexpressed HEK293T cells, were transfected with the 5-′UTR construct. Cells were incubated for 24h after transfection, washed twice with PBS and reconstituted in PBS. An equal volume of the sample and Luciferase substrate solution was taken in the flat-bottom 96-well plate and shaken for 10 minutes before fluorescence reading was measured. To determine the HIV-1 LTR promoter activity, TZM-bl cells harboring LTR-luciferase reporter were reversely transfected with CRISPR Cas13X.1-crRNA for 48 h, followed by TNF treatment for 24 h. Luminescence was measured and normalized to total proteins quantified by BCA assay.

### Cell culture

TZM-bl and J-Lat 10.6 (Cat. #9849), cell lines were obtained from the National Institutes of Health (NIH) AIDS reagent program. The Jurkat-derived CA5 cell line latently infected with replication-competent, full-length HIV-1 genome was provided by Dr. Olaf Kutsch (36,37). TZM-bl, HEK293T (Cat. # CRL-3216, ATCC), and HeLa-MAGI cells were cultured in Dulbecco’s modified Eagle’s medium (DMEM, Cat #D5796, Sigma). THP89GFP HIV-1 latency cell lines were kindly provided by David Levy (New York University). All monocytes and T cell lines were cultured in Roswell Park Memorial Institute (RPMI) 1640 medium (Cat # 11875093, Gibco). Completed cell culture medium contained 10% fetal bovine serum (FBS, Cat. # 10437028, Thermo Fisher), penicillin (100 U/ml)/streptomycin (100 μg/ml) (Cat. # MT30002CI, Corning). Primary peripheral blood mononuclear cells (PBMCs) were maintained and cultured with a completed RPMI medium with 1 × minimum essential medium nonessential amino acid (Cat #11140-050, Gibco), 1 × sodium pyruvate (Cat #11360-070, Gibco), and 20 mM HEPES (Cat #15630-080, Gibco). Human recombinant IL-2 (rIL-2, Roche) at 30 U/ml was supplied to primary cells every 2 days (38), days followed by activation with Cd3/Cd28 for 3 days.

### Flow Cytometry

HIV-1 latency cell line, J-Lat 10.6, containing integrated, GFP reporter– fused viral genomes, were seeded at 5 × 10^5^ cells/ml and transfected with CRISPR-Cas13X.1 crRNA 48 hours followed by treatment with TNF-α (1 ng/ml) for 24 h.Cells were harvested, washed twice with 1× D-PBS, and analyzedusing an Accuri C6 Plus flow cytometer (BD Biosciences) with forwardversus side scatter (FSC-A versus SSC-A) gating. Fluorescence-stained cells were washed three times with 1× Perm/Wash buffer. The staining buffer [1×D-PBS with 2% bovine serum albumin (BSA)] was added to resuspendcells, followed by flow cytometry analysis using the BD Accuri C6 Plus with corresponding optical filters. Fluorescence-stainedcells were analyzed by flow cytometry. The MFI and the percentageof fluorescence-positive cells were determined by using the FlowJoV10 software.

### Quantitave Real-Time PCR

Total RNAs were extracted using the NucleoSpin RNA extraction kit (Macherey-Nagel, Cat#740955.250) following the protocol provided by the manufacturer. RNA samples were reverse transcribed to cDNAs using iScript (BioRad, Cat#1708891). Real-time quantitative PCR (qPCR) was performed on a CFX96 instrument (Bio-Rad), by mixing 5 μl (2×) SYBR Green Supermix (BioRad, Cat#1725214), 0.5 μl primer mix, and 4.5 μl of cDNA template. Data were analyzed by using the Ct method with GAPDH as an internal control.

### RNA immunoprecipitation with qRT-PCR (RIP-qRT-PCR)

To identify the TAR RNA fragments that associate with HIV-1 Tat, RIP assays were performed following the co-transfection of TZM-bl cells with CRISPR Cas13X.1-crRNA and PQCXIP-Flag-Tat. In brief, cells were crosslinked with paraformaldehyde and lysed using NT2 buffer. Cell lysates were incubated with an anti-Flag and anti-Tat antibodies absorbed to the magnetic beads (Thermo Scientific, Cat#88803) for overnight at 4◦C. The DNA-free DNA Removal Kit (Invitrogen, Cat#AM1906) was used to remove DNAs. Extracted RNA samples were subjected to the above qPCR analysis. All qPCR primers were listed in Supplementary Table S1.

### Western blotting

After transfection and infections of cells with HIV-1 IIIB cells were washed with cold PBS and lysed at 4 °C in 1X RIPA lysis buffer (company and cat no), protease inhibitor [company] for 30 mins at 4 °C. Lysates were separated by SDS-PAGE followed by electrophoresis and transfer on to polyvinylidene difluoride (PVDF) membranes which were blocked in 5% nonfat dry milk (NFDM), then incubated with primary antibodies overnight at 4°C followed by washing and incubation with HRP-conjugated secondary antibodies. The blots were developed using the LumiGLO chemiluminescence detection system (Cell Signalling Technology).

### Nested qPCR Assay to Quantify HIV-1 Viral Load

HIV-1 viral load was quantified by nested qPCR, adapted from a previously established protocol (Mousseau et al., 2015). Culture supernatants from HIV-1-infected or reactivated cells were collected, and viral RNA was extracted using the QIAamp Viral RNA Kit (Qiagen) per the manufacturer’s protocol. Extracted RNA was treated with DNase I (Invitrogen) for 10 min at 25°C, followed by enzyme inactivation with EDTA at 65°C for 10 min.A first-round reverse transcription–PCR was then performed using the SuperScript III One-Step RT-PCR System with Platinum Taq High Fidelity (Life Technologies) in a 50-µL reaction, with HIV-1 gag-specific primers O1 and O2. Thermal cycling consisted of reverse transcription at 50°C for 30 min, initial denaturation at 94°C for 2 min, 16 cycles of amplification (94°C, 15 s; 62°C, 30 s; 68°C, 60 s), and a final extension at 65°C for 5 min. The resulting pre-amplified products were purified with the QIAquick PCR Purification Kit (Qiagen) prior to second-round amplification.Nested qPCR was subsequently carried out in a 25-µL reaction using TaqMan Universal PCR Master Mix (Applied Biosystems) with primers O3/O4 and a corresponding TaqMan probe. Cycling parameters included a UNG decontamination step (50°C, 2 min), initial denaturation (95°C, 10 min), and 45 cycles of amplification (95°C, 15 s; 50°C, 20 s; 60°C, 1 min). All reactions were run on a CFX Connect real-time PCR detection system (Bio-Rad). Absolute viral copy number was determined by reference to a standard curve generated from serial dilutions of HIV-1 IIIB stock of known titer.

### Statistical analysis

Flow cytometry data were analyzed using FlowJo v10, and graphs were generated with GraphPad Prism v10.0. Data are presented as mean ± SD. Statistical significance between groups was assessed using un-paired student t-tests, one-way ANOVA, or two-way ANOVA, as appropriate. *P < 0.05, were considered statistically significant.

## RESULTS

### Design and construction of the CRISPR Cas13X.1 system

The Cas13 protein and its CRISPR RNA (crRNA) together form an RNA-guided ribonuclease, which leads to the degradation of the target RNA (39,40). In order to confirm the Cas13X.1 ribonuclease activity, we designed and cloned crRNA targeting mCherry RNA according to previous study (41). Constructed plasmid simultaneously expresses the human-optimized Cas13X.1 protein and a specific crRNA (**Figure 1A**). To analyze the knockdown activity, we transfected HEK293T cells either with the mCherry-targeting or non-targeting (NT) plasmids for 2days. We observed that the relative expression of mCherry mRNA levels significantly decreases in mCherry specific crRNA transfected cells compared to non-targeting (NT) crRNA transfected cells (**Figure 1B**). There are two BbSI restriction sites upstream of the DR sequence. After digesting the plasmid with BbSI restriction enzyme, we observed a band of approximately 8000 bp, which was gel-extracted and proceeded to the next step (**Figure S1A**). In addition fluorescence intensity of the mCherry protein decreased significantly in the mCherry-targeting group (**Figure 1C**). In the mock control group (without transfection), there was no mCherry fluorescence, suggesting that the fluorescence was from the exogenous gene expression. Furthermore, the number of mCherry fluorescence-positive cells analyzed by Flow cytometry was significantly reduced in the mCherry targeting group compared to the non-targeting group (**Figure 1C**, **1D**).

**Figure 1.**
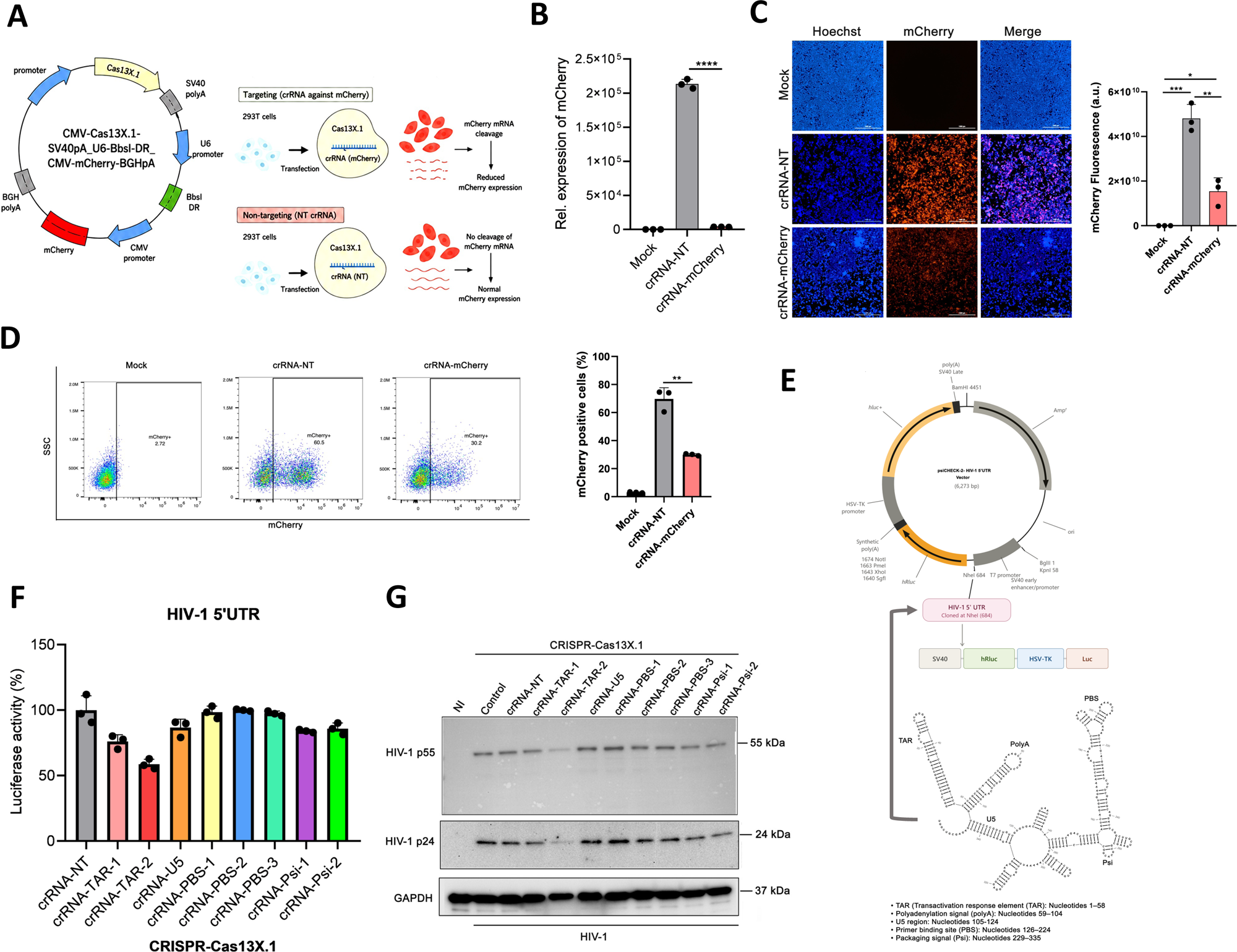
Construction of the CRISPR-Cas13X.1 system to target mCherry expression and the HIV-1 5′ untranslated region (5′UTR). **(A)** Schematic representation of the CRISPR–Cas13X.1 system designed to target mCherry in 293T cells. **(B)** 293T cells were mock transfected or transfected with the CRISPR–Cas13X.1 system containing either a non-targeting crRNA (crRNA-NT) or an mCherry-targeting crRNA (crRNA-mCherry). mCherry RNA levels were quantified by qRT–PCR. Data represent mean ± S.D. (*n* = 3 independent experiments); \*\*\*\**P* < 0.0001. **(C, D)** 293T cells were treated as in (B) and analyzed by immunofluorescence for mCherry expression (scale bar = 1000 µm) (C) and by flow cytometry (CytoFLEX) for quantification of mCherry-positive cells (D). Relative mCherry fluorescence intensity (C) and the percentage of mCherry-positive cells (D) are shown for Mock, crRNA-NT, and crRNA-mCherry conditions. Data represent mean ± S.D. (*n* = 3 independent experiments); \**P* < 0.05, \*\**P* < 0.01, \*\*\**P* < 0.001. **(E)** The secondary structure of the full-length HIV-1 5′ untranslated region (5′ UTR) was predicted using the RNAfold web server and constructed using RNAcanvas, highlighting the cloning site in the psiCHECK-2 reporter vector. **(F)** HEK293T cells were co-transfected with the constructed psiCHECK-2-HIV-1 5′ UTR reporter plasmid and Cas13X.1 crRNA targeting different regions of the HIV-1 5′ UTR or a non-targeting crRNA (crRNA-NT). Dual-luciferase reporter assays were performed, and the Renilla/firefly luciferase activity ratio was calculated to evaluate the effects of targeting different regions of the HIV-1 5′ UTR. Data represent *n* = 3 independent experiments. **(G)** MAGI cells were transfected with Cas13X.1 crRNAs targeting different regions of the HIV-1 5′ UTR or a non-targeting crRNA (crRNA-NT) prior to HIV-1 infection. Cells were subsequently lysed, and HIV-1 p55 and p24 protein levels were analyzed by immunoblotting using an anti-HIV-1 p24 antibody. Equal protein loading was confirmed by immunoblotting with an anti-GAPDH antibody. Data represent n=3 independent experiments.

To test whether CRISPR Cas13X.1 targets the HIV-1 5′UTR, we amplified and cloned the full-length HIV-1 5′UTR, which contains several structural and sequence motifs essential for viral replication, into a luciferase reporter plasmid harbouring both Firefly and Renilla Luciferase genes (psicheck-2) (**Figure 1E**). HEK293T cells were transfected with this construct along with either a crRNA targeting a specific site or a non-targeting crRNA, and luciferase activity was measured according to the instructions from previous studies (42). Luciferase activity was downregulated in the presence of crRNA targeting TAR RNA compared to that of non-targeting crRNA (**Figure 1F**). The effect of Cas13X.1 crRNAs on HIV-1 expression was tested by transfecting MAGI cells with either a crRNA targeting the HIV-1 5′UTR or a non-targeting (NT) crRNA prior to infection with HIV-1. Levels of the Gag precursor p55 and the mature capsid protein p24 were significantly reduced in cells expressing the TAR-2, Psi-1 and Psi-2 crRNAs compared with the non-targeting control (**Figure 1G**). We also observed no difference in cell viability among cells expressing the mCherry-targeting (mC), TAR-targeting, or non-targeting (NT) crRNAs (**Figure S1B**). Therefore, we further explored crRNA targeting of TAR sites, which are Tat-binding sites and sites of transcriptional activation.

### CRISPR-Cas13X.1 crRNA targets the Transactivation response element (TAR) RNA of the HIV-1 5’UTR and disrupts the binding between HIV-1 Tat and TAR

The 5′ untranslated region (5′UTR) of the HIV-1 RNA genome is a structured and highly conserved sequence that regulates diverse processes during the viral replication cycle, including RNA splicing, translation initiation, genome dimerization and packaging into assembling virions, and initiation of reverse transcription (43). The HIV-1 trans-activation response (TAR) element is a highly structured RNA stem-loop located at the 5′ end of all nascent viral transcripts that serves as the essential binding site for the viral Tat protein, enabling recruitment of the host P-TEFb complex and subsequent activation of processive transcription from the HIV-1 LTR (25,30). Given its indispensable role in Tat-dependent transactivation, we sought to determine whether the CRISPR-Cas13X.1 system could be programmed to directly target TAR RNA and thereby disrupt this interaction. The CRISPR-Cas13X.1 plasmid system was constructed to target the HIV-1 5′UTR, specifically the TAR RNA site, as shown in the schematic representation (**Figure 2A**). Among the crRNAs screened, crRNA-TAR-2 produced the most consistent effect, reducing *Renilla* luciferase activity from the HIV-1 5′UTR reporter and lowering HIV-1 p55 and p24 levels in de novo infection. We therefore selected crRNA-TAR-2 for subsequent mechanistic experiments. To confirm its activity in this setting, HEK293T cells were co-transfected with the psiCHECK-2–HIV-1 5′UTR reporter and either crRNA-TAR-2 or crRNA-NT. Luciferase activity was significantly reduced in crRNA-TAR-2–transfected cells relative to crRNA-NT controls (**Figure 2B**).To determine whether Cas13X.1 can be programmed to interfere with Tat-dependent transactivation of the HIV-1 LTR, we co-transfected TZM-bl cells with either crRNA-TAR-2 or crRNA-NT and HIV-1 Tat (pQCXIP-Flag-Tat) to activate HIV-1 LTR-driven transcription. Expression of HIV-Tat was confirmed by immunoblotting with an anti-Flag antibody (**Figure S2A**). Luciferase reporter assays showed that crRNA-TAR-2 transfection decreased HIV-1 LTR activity relative to the crRNA-NT control, indicating that Cas13X.1 targeting of the TAR element attenuates Tat-mediated transactivation (**Figure 2C**).

**Figure 2.**
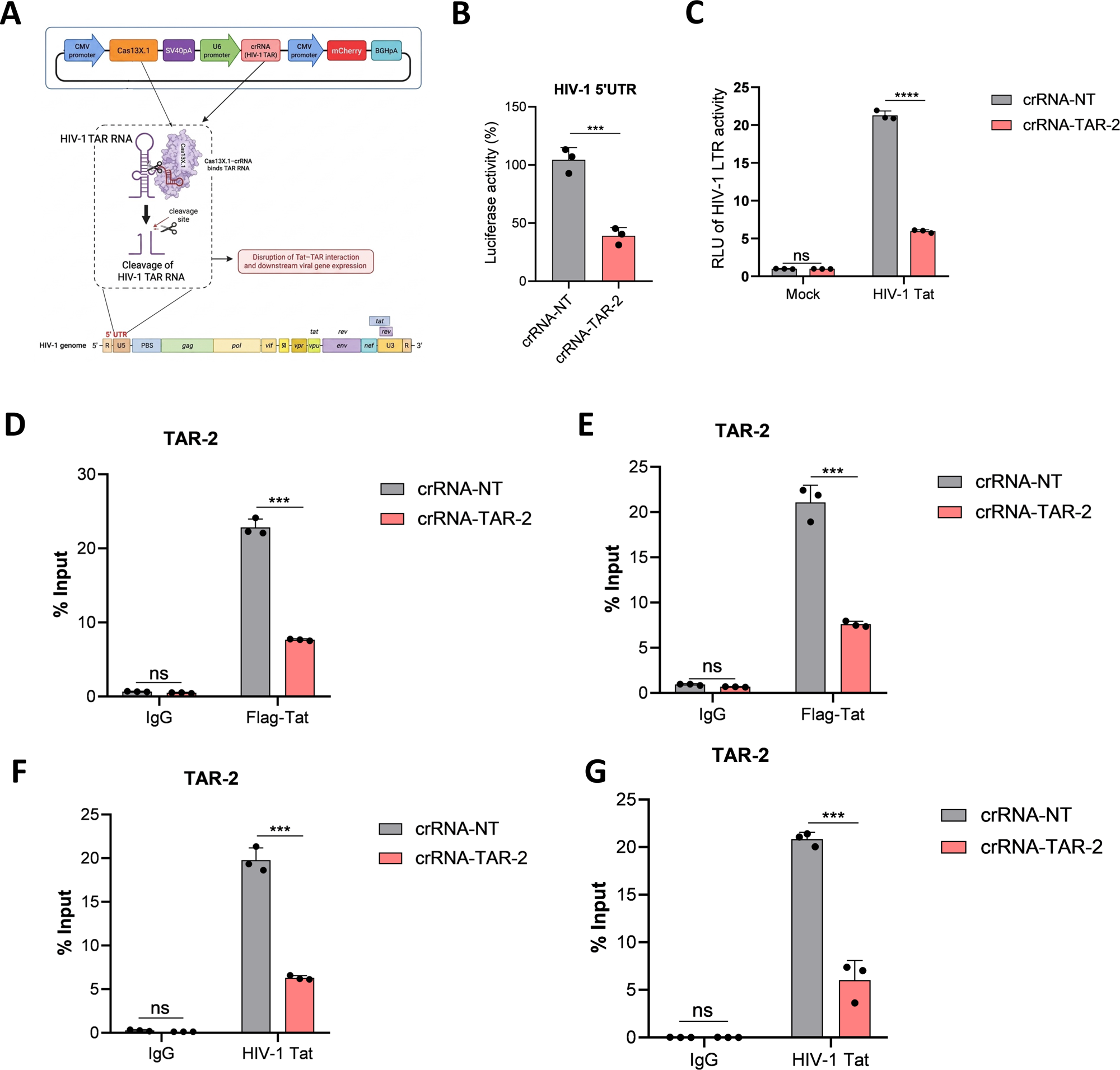
Transactivation response elements (TAR) RNA of HIV-1 5’UTR was targeted using the CRISPR-Cas13X.1 system. **(A)** Schematic representation of HIV-1 5′UTR TAR RNA targeting using the CRISPR-Cas13X.1 crRNA system. **(B)** Luciferase activity was measured following 293T cells were co-transfected with psiCHECK-2-HIV-1 5′UTR reporter plasmid and either Cas13X.1-crRNA-TAR-2 (crRNA-TAR-2) or non-targeting crRNA (crRNA-NT) for 48 h. Data represent mean ± S.D. (*n* = 3 independent experiments); \*\*\**P* < 0.001. **(C)** TZM-bl cells were co-transfected with the Cas13X.1 system (crRNA-TAR-2 or crRNA-NT) and pQCXIP-Flag-Tat plasmid or empty vector (Vector). HIV-1 LTR activity was assessed by measuring luciferase activity in co-transfected samples. Data represent mean ± S.D. (*n* = 3 independent experiments); ns, not significant; \*\*\**P* < 0.001. **(D, E, F, G)** TZM-bl cells were co-transfected with pQCXIP-Flag-Tat and Cas13X.1 crRNA-NT or crRNA-TAR-2 for 48 h, followed by RNA immunoprecipitation (RIP) with anti-Flag or IgG (control) antibody (D, E) and with anti-Tat or IgG (control) antibody (F, G). RIP-qRT-PCR analysis was performed using two independent TAR-specific primer pairs, to confirm Tat–TAR RNA association. Data represent mean ± S.D. (*n* = 3 independent experiments); ns, not significant; \*\*\**P* < 0.001.

To evaluate the mechanistic basis of Cas13X.1-mediated disruption of the Tat–TAR interaction, TZM-bl cells were co-transfected with the Tat expression plasmid and either crRNA-TAR-2 or crRNA-NT. Forty-eight hours after transfection, RNA immunoprecipitation (RIP) was performed with an anti-Flag antibody (**Figure 2D, 2E**) or an anti-Tat antibody (**Figure 2F, 2G**), and recovered RNA was quantified by RT-qPCR using two independent primer sets spanning the TAR-1 and TAR-2 target sites. TAR RNA recovery, expressed as percent input, was significantly reduced in crRNA-TAR-2–transfected cells relative to crRNA-NT controls with both antibodies and both primer sets (**Figure 2D–2G**), indicating that Cas13X.1 targeting of TAR impairs the Tat–TAR RNA interaction.

### HIV-1 TAR RNA targeted by the CRISPR-Cas13X.1 system suppresses viral gene expression in de novo infections

To determine whether a Cas13X.1 crRNA directed against the HIV-1 5′ untranslated region (5′UTR), specifically the trans-activation response (TAR) element, could restrict viral replication, we transfected MAGI cells with either a TAR-targeting crRNA (crRNA-TAR-2) or a non-targeting control crRNA (crRNA-NT) and subsequently challenged the cells with HIV-1 for 48 h. Cells expressing TAR crRNA exhibited markedly reduced levels of the Gag precursor p55 and the mature capsid protein p24, relative to NT crRNA-transfected controls (**Figure 3A**), indicating suppression of viral protein production. Consistent with this reduction, quantification of viral transcripts by RT-qPCR revealed decreased steady-state levels of both *gag* and *tat* mRNA in crRNA-TAR-2 transfected cells compared with crRNA-NT controls (**Figure 3B, 3C**). Immunofluorescence staining for HIV-1 p24 corroborated these findings, showing a visibly reduced proportion of p24-positive cells in the crRNA-TAR-2 condition relative to the crRNA-NT condition (**Figure 3D**).

**Figure 3.**
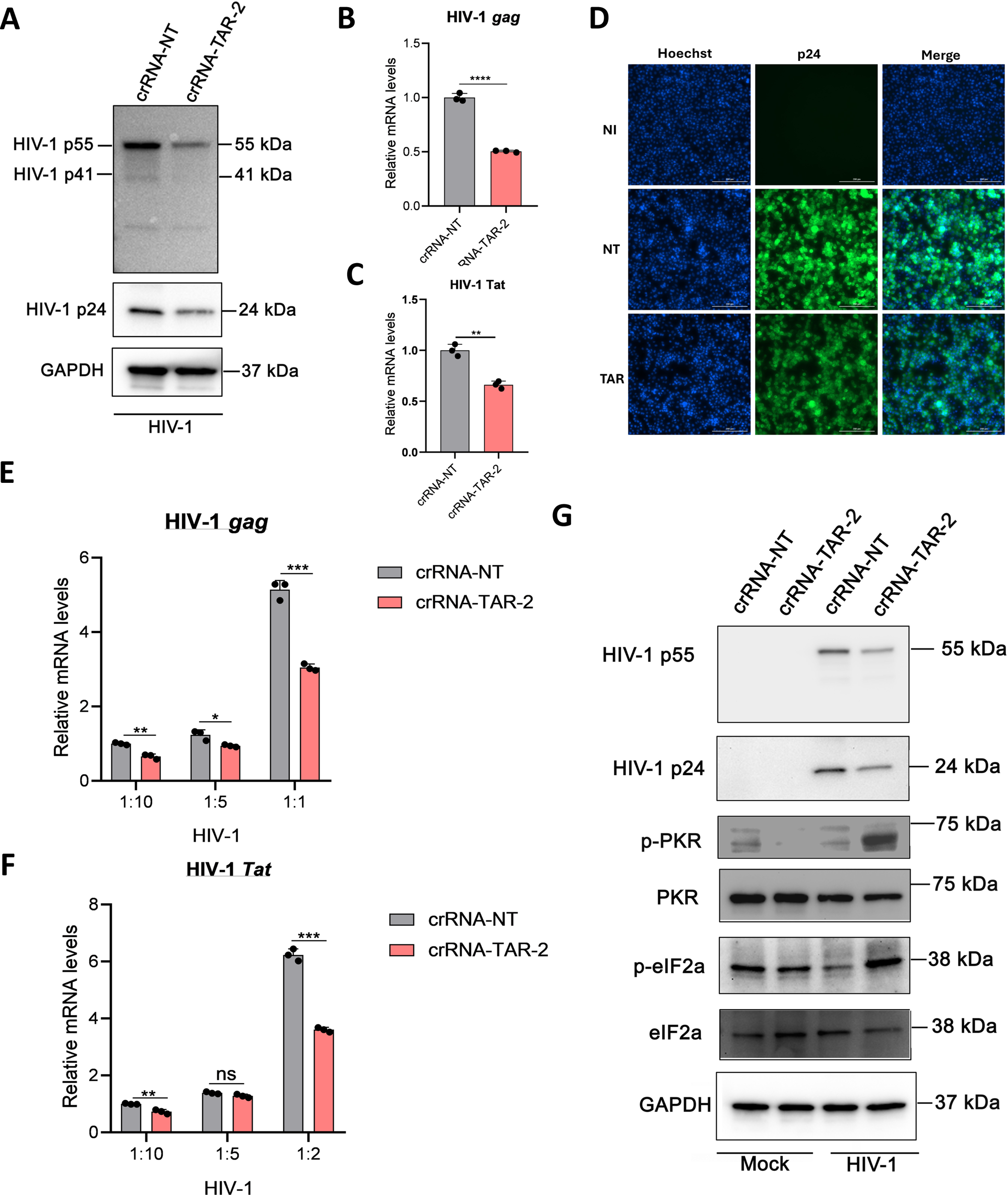
Cas13X.1 crRNA targeting HIV-1 TAR RNA suppresses de novo infection. (A–C) MAGI cells were transfected with the Cas13X.1 system containing non-targeting crRNA (crRNA-NT) or TAR-targeting crRNA (crRNA-TAR-2) prior to infection with HIV-1 IIIB. **(A)** Immunoblotting of cell lysates using anti-HIV-1 p24 antibody; equal loading was confirmed by reprobing with anti-GAPDH antibody. Blots are representative of three independent experiments. **(B, C)** HIV-1 *gag* (B) and *tat* (C) transcript levels were analyzed by qRT-PCR. Data represent mean ± S.D. (*n* = 3 independent experiments); \*\**P* < 0.01, \*\*\**P* < 0.001. **(D)** MAGI cells were treated as in (A–C) and subjected to immunofluorescence staining with anti-HIV-1 p24 antibody. Images are representative of three independent experiments. Scale bar = 200 µm. **(E, F)** MAGI cells were infected with serial dilutions (1:1, 1:5, 1:10) of viral supernatant collected from the infected cultures in (A-C), and HIV-1 *gag* (E) and *tat* (F) mRNA levels were analyzed by qRT-PCR. Data represent mean ± S.D. (*n* = 3 independent experiments); ns, not significant; \**P* < 0.05, \*\**P* < 0.01, \*\*\**P* < 0.001. **(G)** MAGI cells were transfected with the Cas13X.1 system containing crRNA-NT or crRNA-TAR-2, followed by HIV-1 infection or left uninfected (Mock), and subjected to immunoblotting for HIV-1 p24, p-PKR, total PKR, p-eIF2α and total eIF2α. Equal protein loading was confirmed by reprobing the blot with anti-GAPDH antibody.

To determine whether TAR-targeted knockdown was sustained across sequential rounds of infection, we performed a second-round (“part II”) infection assay in which cell-free supernatant from the initial infection was serially diluted and used to inoculate freshly plated cells. RT-qPCR analysis of this second-round infection again demonstrated significantly reduced *tat* and *gag* transcript levels in supernatant derived from the crRNA-TAR-2 condition relative to the crRNA-NT condition across the dilution series (**Figure 3E, 3F**), indicating that Cas13X.1-mediated targeting of TAR RNA durably restricts progeny virus production and downstream infectivity.

In addition, TAR RNA can either activate or inhibit the double-stranded RNA-dependent protein kinase (PKR), depending on its concentration. Activation of PKR through autophosphorylation triggers an antiviral response by promoting phosphorylation of the α-subunit of eukaryotic initiation factor 2 (eIF2α), thereby inhibiting translation initiation and reducing viral protein synthesis (43–46). In contrast, inhibition of PKR activity, accompanied by reduced eIF2α phosphorylation, can relieve this translational restriction and consequently promote viral protein synthesis (44,45,47). We observed increased expression of phosphorylated PKR (p-PKR) and phosphorylated eIF2α (p-eIF2α) in crRNA-TAR-2-transfected cells compared to crRNA-NT-transfected cells following HIV-1 infection (**Figure 3G**). In the absence of infection, transfection with the Cas13X.1 system alone induced modest p-eIF2α activation in both crRNA-TAR-2 and crRNA-NT conditions, with no significant difference between the two. These findings suggest that TAR-targeting by Cas13X.1 enhances activation of the PKR–eIF2α antiviral response during HIV-1 infection.

### HIV-1 TAR-directed Cas13X.1 crRNA limits latency reactivation and suppresses viral load and gene expression in PBMCs from HIV-1-positive individuals

We next asked whether TAR-directed Cas13X.1 crRNA could similarly restrict viral gene expression in the context of HIV-1 latency. In the J-Lat 10.6 latency model, stimulation with tumor necrosis factor-α (TNF-α) induced the expected increase in the proportion of GFP-positive cells, reflecting reactivation of the integrated provirus. Transfection with crRNA-TAR-2, prior to TNF-α stimulation, significantly reduced the frequency of GFP-positive cells relative to crRNA-NT transfected controls (**Figure 4A**), demonstrating that Cas13X.1-mediated TAR targeting attenuates reactivation-associated viral gene expression from latent reservoirs.

**Figure 4.**
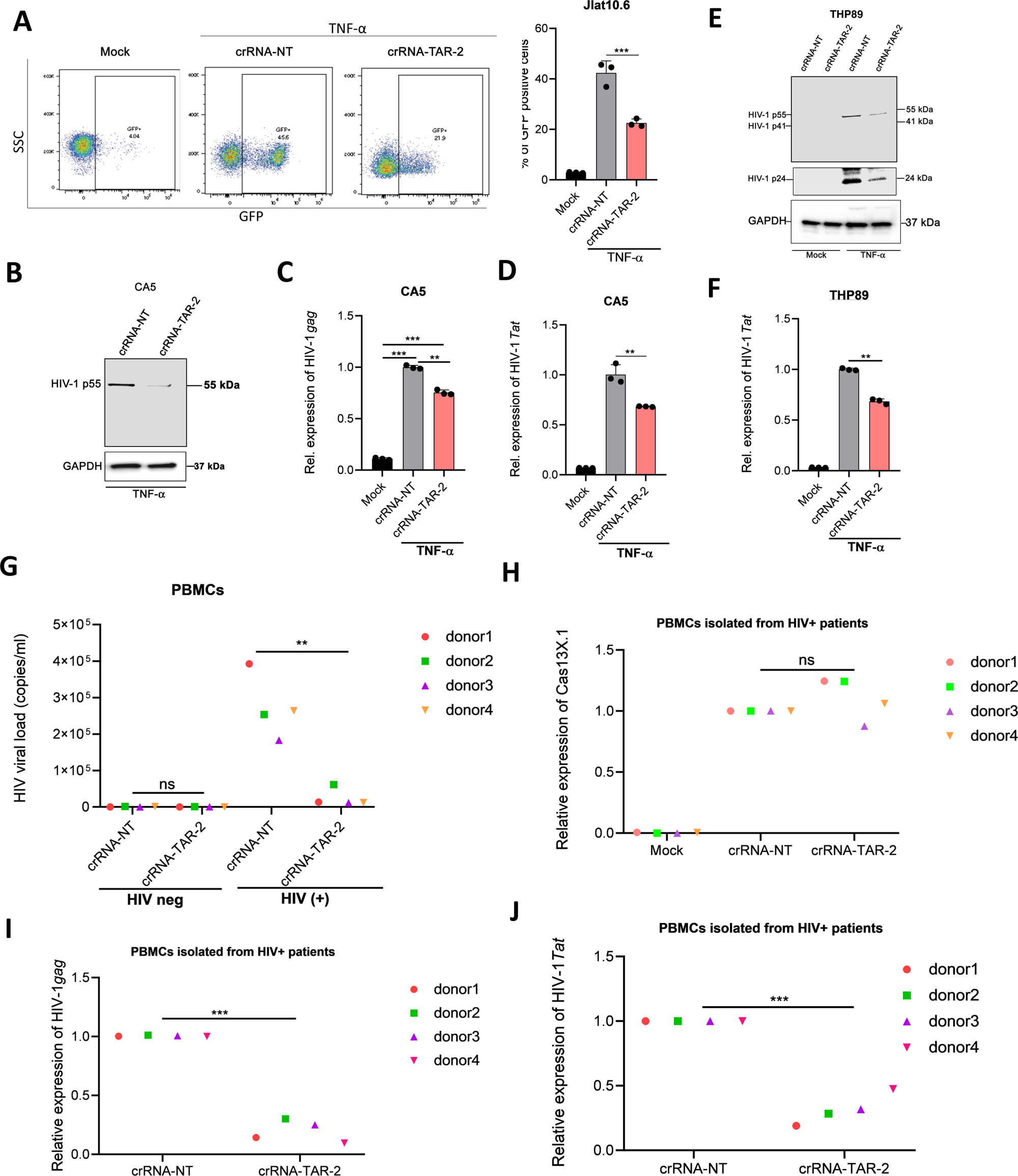
CRISPR-Cas13X.1 crRNA targets HIV-1 TAR RNA in a latency infection model and suppresses viral replication in PBMCs from HIV-1-positive patients. (A) J-Lat 10.6 cells were left untransfected (Mock) or transfected with the Cas13X.1 system containing non-targeting crRNA (crRNA-NT) or TAR-targeting crRNA (crRNA-TAR-2), followed by treatment with TNF-α (1 ng/mL) for 24 h. Cells were analyzed by flow cytometry (CytoFLEX) for the percentage of GFP-positive cells. Data represent mean ± S.D. (*n* = 3 independent experiments); \*\**P* < 0.01. **(B–D)** CA5 cells were left untransfected (Mock) or transfected with the Cas13X.1 system containing crRNA-NT or crRNA-TAR-2, followed by treatment with TNF-α (1 ng/mL) to activate GFP expression. **(B)** Western blot analysis of cell lysates using anti-HIV-1 p24 antibody, comparing crRNA-NT and crRNA-TAR-2–transfected cells. Equal loading was confirmed by reprobing with anti-GAPDH antibody. **(C, D)** HIV-1 *gag* (C) and *tat* (D) transcript levels were analyzed by qRT-PCR. Data represent mean ± S.D. (*n* = 3 independent experiments); \*\**P* < 0.01, \*\*\**P* < 0.001. **(E, F)** THP89 cells were transfected with the Cas13X.1 system containing crRNA-NT or crRNA-TAR-2, followed by treatment with TNF-α (1 ng/mL) or left untreated (Mock) for 24 h. **(E)** Cell lysates were immunoblotted with anti-HIV-1 p24 antibody; equal loading was confirmed by reprobing with anti-GAPDH antibody. **(F)** HIV-1 *tat* transcript levels were analyzed by qRT-PCR. Data represent mean ± S.D. (*n* = 3 independent experiments); \*\**P* < 0.01. **(G)** Peripheral blood mononuclear cells (PBMCs) were isolated from HIV-1–negative (*n* = 4) and HIV-1–positive (*n* = 4) donors, activated with anti-CD3/CD28 antibodies for 3 days, and transfected with the Cas13X.1 system containing non-targeting crRNA (crRNA-NT) or TAR-targeting crRNA (crRNA-TAR-2). Cell-free HIV-1 viral load was quantified by two-step nested qPCR, referenced against a known copy number standard derived from HIV-1 IIIB stock titered using the Takara viral titer kit. Data represent mean ± S.D (n= 4 independent donors).; ns, not significant ; \*\**P* < 0.01 **(H–J)** PBMCs from HIV-1-positive donors (*n* = 4) were activated as in (A) and transfected with the Cas13X.1 system containing crRNA-NT or crRNA-TAR, or left untransfected (Mock). Cell lysates were analyzed by qRT-PCR for Cas13X.1 (H), HIV-1 *gag* (I), and HIV-1 *tat* (J) mRNA levels. Data represent mean ± S.D. (*n* = 4 independent donors); ns, not significant; \*\*\**P* < 0.001.

We extended this analysis to the CA5 cell line, an additional CD4⁺ T cell–based model of HIV-1 latency that, similar to J-Lat clones, harbours a stably integrated, transcriptionally silent provirus reactivatable by TNF-α stimulation (48). In this model, crRNA-TAR-2 transfection similarly reduced Gag precursor HIV-1 p55 protein levels and *gag* and *tat* transcript levels relative to crRNA-NT controls (**Figure 4B–D**), confirming the knockdown effect across an independent latency model. Furthermore, we evaluated THP89 cells, a monocytic latency model in which TNF-α stimulation reactivates provirus. Transfection with crRNA-TAR-2 reduced Gag precursor HIV-1 p55 and viral capsid protein HIV-1 p24 levels and decreased tat mRNA expression relative to crRNA-NT–transfected controls (**Figure 4E, 4F**), indicating that TAR-directed knockdown also suppresses reactivated HIV-1 gene expression in myeloid-lineage latently infected cells.

To extend these findings beyond cell line models, we next evaluated the efficacy of the Cas13X.1–TAR-2 crRNA system in primary peripheral blood mononuclear cells (PBMCs) obtained from HIV-1-positive and HIV-1-negative donors (n = 4 donors per group). PBMCs were activated with anti-CD3/CD28 stimulation for 3 days and subsequently transfected with either TAR-targeting or NT control crRNA. Quantification of cell-free HIV-1 viral load in culture supernatants revealed a significant reduction in crRNA-TAR-2 transfected PBMCs relative to crRNA-NT transfected PBMCs from HIV-1-positive donors, as expected, viral load remained undetectable in PBMCs from HIV-1-negative donors (**Figure 4G**). To confirm efficient delivery of the Cas13X.1 system, we measured Cas13X.1 transcript levels by RT-qPCR in HIV-1-positive PBMCs. Cas13X.1 mRNA was significantly enriched in Cas13X.1 crRNA-transfected cells relative to untransfected (Mock) controls, with comparable expression levels between crRNA-NT and crRNA-TAR-2 transfected cells (**Figure 4H**), confirming that differences in viral outcome were attributable to target-specific knockdown rather than differential transfection efficiency. Consistent with the reduced viral load, crRNA-TAR-2-transfected PBMCs from HIV-1-positive donors showed significantly lower gag and tat mRNA levels than crRNA-NT-transfected controls (Figure 4I and 4J). Together, these results demonstrate that Cas13X.1-mediated targeting of the HIV-1 TAR element effectively suppresses viral replication and gene expression in primary patient-derived PBMCs, supporting the translational potential of this approach.

## DISCUSSION

The 5′ untranslated region (5′UTR) of the HIV-1 genome is the most highly conserved region across viral clades and circulating strains (7,49–51) . Despite this compact size, the 5′UTR is structurally and functionally dense, folding into a series of stem-loop RNA elements including the trans-activation response (TAR) element, the polyadenylation (polyA) signal, the primer-binding site (PBS), the dimerization initiation site (DIS), the major splice donor (SD) site, and the packaging signal (Ψ), each of which governs a distinct step of the viral life cycle (7,50,52). Beyond its established role in genome dimerization and packaging, the 5′UTR coordinates transcriptional activation, alternative splicing, and initiation of reverse transcription via primer annealing at the PBS (52). For several retroviruses, elements required for genomic RNA packaging are located downstream of the major 5′ splice donor site, providing a mechanism by which the full-length, unspliced genome is selectively packaged into virions while spliced viral mRNAs are excluded (53). Given the HIV-1 5′UTR’s essentiality, structural conservation, and involvement in multiple steps of the viral life cycle including transcription, splicing, dimerization, and packaging the 5′UTR constitutes a rational target for nucleic acid-based antivirals. Disruption of a single conserved structural element within this region could simultaneously impair several of these processes, and the accompanying sequence constraints may limit the mutational routes available for escape, potentially yielding a higher genetic barrier to resistance than strategies aimed at more variable coding regions (49,54,55).

Among the structural elements within the HIV-1 5′ untranslated region (5’ UTR), the transactivation response (TAR) RNA stem-loop is the most extensively characterized, serving as the binding platform through which the viral Tat protein activates transcription. Positioned at the 5′ terminus of all HIV-1 transcripts, TAR adopts a conserved RNA hairpin containing a trinucleotide pyrimidine bulge and an apical loop, structural features that together confer high-affinity Tat binding and enable recruitment of the host P-TEFb elongation complex as part of the larger super elongation complex (SEC), a step indispensable for proviral transactivation (25–30). This Tat-TAR interaction is essential not only for active viral replication but also for reactivation of HIV-1 from latency, making it a long-pursued target for the development of antiviral and latency-reversing therapeutic strategies (56).

Beyond its role in Tat-dependent transactivation, TAR RNA engages a broad network of host proteins that shape both gene expression and antiviral signaling, including TAR RNA-binding protein (TRBP), protein kinase R (PKR), RNA helicase A, negative elongation factor (NELF), nuclear factor 90 (NF90), and La antigen (47,57–61). Among these, PKR is of particular importance as a central effector of the interferon-mediated antiviral response, its latent kinase activity is triggered by double-stranded RNA structures such as those formed during viral replication or bidirectional transcription, and once activated, PKR autophosphorylates and in turn phosphorylates eukaryotic translation initiation factor 2α (eIF2α), shutting off protein synthesis and triggering apoptotic pathways as part of a broader antiviral and antiproliferative program (62–64). HIV-1 counteracts this defense by exploiting the structural properties of TAR itself, TAR’s stem-loop architecture and its association with TRBP uncouple PKR binding from PKR activation, allowing the virus to sequester and functionally suppress PKR without triggering its autophosphorylation, thereby keeping eIF2α unphosphorylated and preserving viral protein synthesis (62,65,66). In our study, targeting TAR RNA with CRISPR-Cas13X.1 was associated with increased phosphorylation of both PKR and eIF2α during HIV-1 infection compared with the non-targeting crRNA control. This enhanced activation of the PKR–eIF2α pathway may contribute to the reduction in viral gene expression observed following TAR targeting, suggesting that Cas13X.1-mediated disruption of TAR RNA may not only interfere with Tat-dependent transcription but may also favor activation of an intrinsic antiviral response that restricts viral protein synthesis.

Unlike the DNA-targeting CRISPR-Cas9 system, which introduces double-strand breaks and permanent modifications into the host or proviral genome (67), the Cas13 family of class 2, type VI CRISPR-Cas effectors are RNA-guided, RNA-targeting ribonucleases that recognize and cleave single-stranded RNA without altering the underlying DNA sequence (14). All-in-one AAV plasmids co-packaging Cas13bt3 (also known as Cas13X) with a single VEGFA-targeting sgRNA have been shown to achieve potent VEGFA knockdown across multiple model systems, including human cell lines, 3D human embryonic stem cell (hESC)-derived retinal organoids, and humanized VEGF-transgenic mouse retinae in vivo (68).

Cas13X.1, identified from metagenomic mining of hypersaline environmental samples, is a markedly compact effector of only 775 amino acids, among the smallest Cas13 orthologs characterized to date, while retaining robust RNA knockdown activity in mammalian cells and tolerance for single-nucleotide mismatches in target recognition, a property that facilitates its use in prophylactic RNA virus inhibition (16). Unlike other Cas13 proteins, Cas13X.1 does not require a protospacer flanking site (PFS) on the target RNA. Furthermore, a single direct repeat (DR) at the 3′ end of the crRNA is sufficient for target recognition and cleavage activity (16). This distinction is particularly relevant for an RNA virus like HIV-1, and for targeting viral transcripts without risking off-target genomic editing or permanent modification of the host cell genome, an important safety consideration for any eventual therapeutic application.

We first established the functionality of the Cas13X.1 platform using a single-vector system (CMV-Cas13X.1-SV40pA_U6-BbsI-DR_CMV-mCherry-BGHpA; Addgene #171379) in which a crRNA targeting mCherry was used to confirm robust, guide-dependent knockdown of a fluorescent reporter transcript by immunofluorescence, flow cytometry and RT-qPCR, confirming on-target cleavage activity prior to application against HIV-1 transcripts. Given the high conservation of the HIV-1 5′UTR across viral strains, we designed a panel of crRNAs tiling the TAR, U5, PBS, and Ψ elements and validated their activity using a cloned HIV-1 5′UTR fused to *Renilla* luciferase in the psiCHECK-2 dual-reporter vector. Among the candidate crRNAs, only the TAR-targeting crRNA produced consistent knockdown of reporter activity, consistent with the essential and structurally accessible nature of the TAR stem-loop, and this crRNA was carried forward for all subsequent functional studies. Because TAR RNA is obligately required for Tat-dependent transcriptional elongation from the HIV-1 LTR, we reasoned that Cas13X.1-mediated cleavage of TAR would not only reduce TAR transcript abundance but also directly interfere with the Tat-TAR ribonucleoprotein interaction that drives processive viral transcription. Using TZM-bl cells, we confirmed that Cas13X.1-crRNA-TAR disrupted the physical interaction between Tat and TAR RNA, as measured by RNA immunoprecipitation (RIP), providing a direct mechanistic link between TAR cleavage and the loss of Tat-dependent transactivation observed downstream. Building on this mechanistic validation, we then demonstrated that Cas13X.1 crRNA-TAR suppresses HIV-1 p24 protein and *tat and gag* transcript levels in de novo-infected MAGI cells, restricts progeny virus production upon second-round infection, and attenuates reactivation-associated viral gene expression across three independent, lineage-distinct models of HIV-1 latency (J-Lat 10.6, CA5, and THP89). Following HIV-1 infection in MAGI cells, phosphorylated PKR (p-PKR) and phosphorylated eIF2α (p-eIF2α) were both upregulated in crRNA-TAR-2–transfected cells relative to crRNA-NT–transfected controls, suggesting that Cas13X.1-mediated TAR disruption enhances antiviral signaling through the PKR-eIF2α pathway. Consistent with this, HIV-1 infection increased p-PKR only modestly while reducing p-eIF2α in crRNA-NT– transfected samples, indicating that the virus limits full activation of the PKR-eIF2α pathway through TAR RNA. Finally, extending this system to primary PBMCs from HIV-1-positive donors, we observed a significant reduction in cell-free viral load and viral transcript levels following crRNA-TAR-2 transfection, with comparable Cas13X.1 delivery efficiency between TAR-2 and NT crRNA conditions, arguing that the antiviral effect reflects target-specific RNA knockdown rather than differences in transfection or expression efficiency. Together, these results indicate that a single, rationally selected crRNA targeting a conserved element of the HIV-1 5′UTR is sufficient to durably suppress viral gene expression across de novo infection, multiple models of latency, and primary patient-derived cells.

In this study, we designed and functionally characterized a CRISPR-Cas13X.1 crRNA targeting the TAR element of the HIV-1 5′UTR, demonstrating that it disrupts the Tat-TAR interaction, restores PKR-eIF2α–mediated antiviral signaling, and suppresses HIV-1 gene expression during de novo infection, latency reactivation across lymphocytic and monocytic models, and in primary PBMCs from HIV-1-positive donors. These findings establish that targeting a single, conserved regulatory RNA element with Cas13X.1 can both directly restrict the viral life cycle and reactivate innate antiviral defenses, offering a broadly effective strategy against HIV-1 across the biological contexts most relevant to a functional cure. Given its compact size relative to other Cas13 orthologs, Cas13X.1 is well suited for single-vector AAV packaging, supporting a plausible path toward durable, gene-therapy-based suppression of HIV-1 replication and reactivation. Future work incorporating stable or viral-vector delivery, primary resting CD4⁺ T cell and humanized mouse models, and comprehensive off-target profiling will be essential next steps toward clinical translation.

## ACKNOWLEDGMENTS

This study was supported by National Institute of Health (NIH) research grants R01DA059538 and R01MH134402 to J.Z.

## SUPPLEMENTARY FIGURES

**Figure S1.**
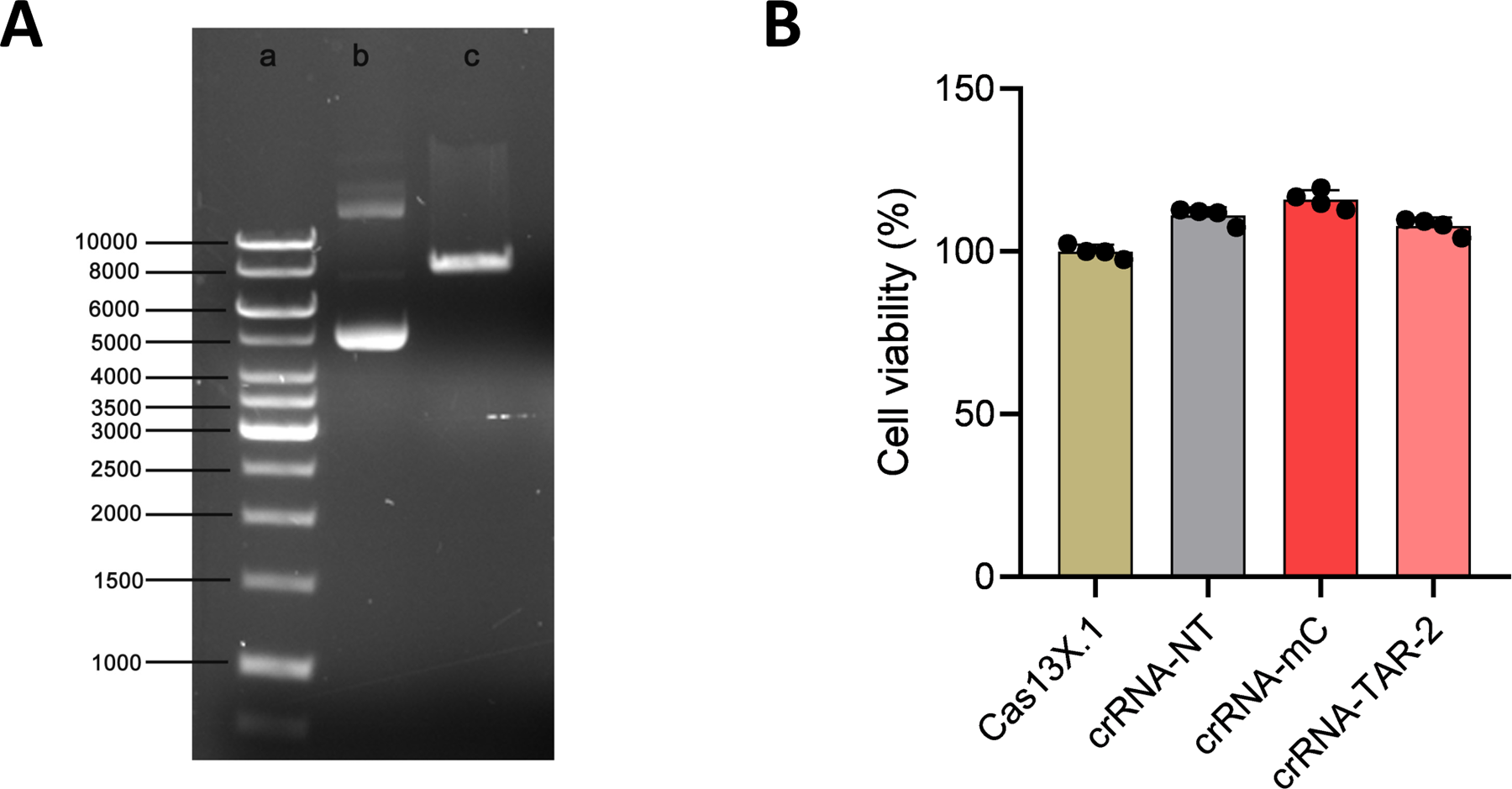
**(A)** The CMV-Cas13X.1-U6-crRNA plasmid was digested with restriction endonuclease BbsI and analyzed by 1% agarose gel electrophoresis. Lane a, 1-kb DNA ladder (GeneRuler); lane b, uncut plasmid; lane c, BbsI-digested plasmid. **(B)** HEK293T cells were transfected with the Cas13X.1 plasmid and the indicated crRNAs (crRNA-NT, crRNA-mCherry, or crRNA-TAR-2) for 48 h. Cell viability was analyzed using the CellTiter-Glo Luminescent Cell Viability Assay.

**Figure S2.**
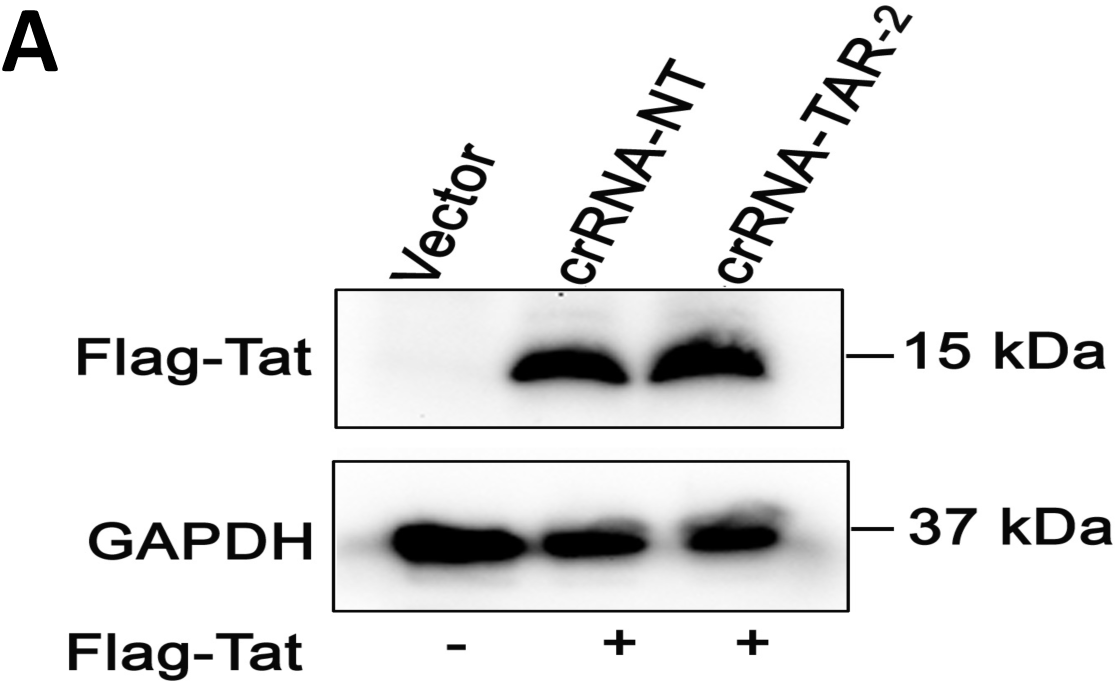
**(A)** Western blot confirming Flag-Tat expression in pQCXIP-Flag-Tat–transfected cells relative to vector-only controls; equal loading was confirmed by reprobing with anti-GAPDH antibody.

## Notes

### Competing Interest Statement

The authors have declared no competing interest.

## REFERENCES

1. Hu WS and Hughes SH: HIV-1 reverse transcription. Cold Spring Harb Perspect Med 2, 2012.

2. Craigie R and Bushman FD: HIV DNA integration. Cold Spring Harb Perspect Med 2, 2012.

3. Ruelas DS and Greene WC: XAn integrated overview of HIV-1 latency. Cell 155: 519, 2013.

4. Churchill MJ, Deeks SG, Margolis DM, Siliciano RF and Swanstrom R: HIV reservoirs: What, where and how to target them. Nat Rev Microbiol 14: 55–60, 2015.

5. Clavel F and Hance AJ: HIV Drug Resistance., 2004.

6. Hayashi T, Jean M, Huang H, Simpson S, Santoso NG and Zhu J: Screening of an FDA-approved compound library identifies levosimendan as a novel anti-HIV-1 agent that inhibits viral transcription. Antiviral Res 146: 76–85, 2017.

7. Jones CP, Cantara WA, Olson ED and Musier-Forsyth K: Small-angle X-ray scattering-derived structure of the HIV-1 5’ UTR reveals 3D tRNA mimicry. Proc Natl Acad Sci U S A 111: 3395–3400, 2014.

8. Cho SW, Kim S, Kim JM and Kim JS: Targeted genome engineering in human cells with the Cas9 RNA-guided endonuclease. Nat Biotechnol 31: 230–232, 2013.

9. Jackson AL, Bartz SR, Schelter J, et al.: Expression profiling reveals off-target gene regulation by RNAi. Nat Biotechnol 21: 635–637, 2003.

10. Herskovitz J, Hasan M, Patel M, et al.: CRISPR-Cas9 Mediated Exonic Disruption for HIV-1 Elimination. EBioMedicine 73: 103678, 2021.

11. Kosicki M, Tomberg K and Bradley A: Repair of double-strand breaks induced by CRISPR–Cas9 leads to large deletions and complex rearrangements. Nat Biotechnol 36: 765–771, 2018.

12. Wang H, La Russa M and Qi LS: CRISPR/Cas9 in Genome Editing and Beyond. Annu Rev Biochem 85: 227–264, 2016.

13. Yoder KE and Bundschuh R: Host Double Strand Break Repair Generates HIV-1 Strains Resistant to CRISPR/Cas9. Sci Rep 6: 29530, 2016.

14. Abudayyeh OO, Gootenberg JS, Konermann S, et al.: C2c2 is a single-component programmable RNA-guided RNA-targeting CRISPR effector. Science (1979) 353, 2016.

15. Makarova KS, Wolf YI, Iranzo J, et al.: Evolutionary classification of CRISPR–Cas systems: a burst of class 2 and derived variants. Nat Rev Microbiol 18: 67–83, 2020.

16. Xu C, Zhou Y, Xiao Q, et al.: Programmable RNA editing with compact CRISPR– Cas13 systems from uncultivated microbes. Nat Methods 18: 499–506, 2021.

17. O’Connell MR: Molecular Mechanisms of RNA Targeting by Cas13-containing Type VI CRISPR–Cas Systems. J Mol Biol 431: 66–87, 2019.

18. Wang D, Zhang F and Gao G: CRISPR-Based Therapeutic Genome Editing: Strategies and In Vivo Delivery by AAV Vectors. Cell 181: 136–150, 2020.

19. Li Y, Xu J, Guo X, et al.: Collateral cleavage of 28s rRNA by RfxCas13d causes death of mice., 2022.

20. Shi P, Murphy MR, Aparicio AO, et al.: RNA-guided cell targeting with CRISPR/RfxCas13d collateral activity in human cells., 2021.

21. Bot JF, van der Oost J and Geijsen N: The double life of CRISPR–Cas13. Curr Opin Biotechnol 78: 102789, 2022.

22. Hu Y, Chen Y, Xu J, et al.: Metagenomic discovery of novel CRISPR-Cas13 systems. Cell Discov 8, 2022.

23. Lu K, Heng X and Summers MF: Structural determinants and mechanism of HIV-1 genome packaging. J Mol Biol 410: 609–633, 2011.

24 . Berkhout B: Structure and Function of the Human Immunodeficiency Virus Leader RNA. pp1–34, 1996.

25. Puglisi JD, Tan R, Calnan BJ, Frankel AD and Williamson JR: Conformation of the TAR RNA-Arginine Complex by NMR Spectroscopy. Science (1979) 257: 76–80, 1992.

26. Aboul-Ela F, Karn J and Varani G: The Structure of the Human Immunodeficiency Virus Type-1 TAR RNA Reveals Principles of RNA Recognition by Tat Protein., 1995.

27. Aboul-Ela F, Karn J and Varani G: Structure of HIV-1 TAR RNA in the absence of ligands reveals a novel conformation of the trinucleotide bulge., 1996.

28. Ippolito JA and Steitz TA: A 1.3-Å resolution crystal structure of the HIV-1 trans-activation response region RNA stem reveals a metal ion-dependent bulge conformation., 1998.

29. Klaver B, Berkhout B and Bosch L: Evolution of a disrupted TAR RNA hairpin structure in the HIV-1 virus ICorresponding author Communicated., 1994.

30. Dingwall C, Ernberg I, Gait MJ, et al.: Human immunodeficiency virus 1 tat protein binds trans-activation-responsive region (TAR) RNA in vitro (trans-activation/RNA-binding proteins/gene synthesis/andsense RNA)., 1989.

31. Berkhout B: Structural features in TAR RNA of human and simian immunodeficiency viruses: a phylogenetic analysis.

32. Yin L, Zhao F, Sun H, et al.: CRISPR-Cas13a Inhibits HIV-1 Infection. Mol Ther Nucleic Acids 21: 147–155, 2020.

33. Nguyen H, Wilson H, Jayakumar S, Kulkarni V and Kulkarni S: Efficient inhibition of hiv using crispr/cas13d nuclease system. Viruses 13, 2021.

34. Zhu J, Gaiha GD, John SP, et al.: Reactivation of Latent HIV-1 by Inhibition of BRD4. Cell Rep 2: 807–816, 2012.

35. Sanjana NE, Shalem O and Zhang F: Improved vectors and genome-wide libraries for CRISPR screening. Nat Methods 11: 783–784, 2014.

36. Kutsch O, Benveniste EN, Shaw GM and Levy DN: Direct and Quantitative Single-Cell Analysis of Human Immunodeficiency Virus Type 1 Reactivation from Latency. J Virol 76: 8776–8786, 2002.

37. Duverger A, Jones J, May J, Bibollet-Ruche F, Wagner FA, Cron RQ and Kutsch O: Determinants of the Establishment of Human Immunodeficiency Virus Type 1 Latency. J Virol 83: 3078–3093, 2009.

38. Zhou D, Hayashi T, Jean M, et al.: Inhibition of Polo-like kinase 1 (PLK1) facilitates the elimination of HIV-1 viral reservoirs in CD4 + T cells ex vivo., 2020.

39. Yoon PH, Zhang Z, Loi KJ, et al.: Structure Guided Discovery of Ancestral CRISPR-Cas13 Ribonucleases.

40. Smargon AA, Cox DBT, Pyzocha NK, et al.: Cas13b Is a Type VI-B CRISPR-Associated RNA-Guided RNase Differentially Regulated by Accessory Proteins Csx27 and Csx28. Mol Cell 65: 618–630.e7, 2017.

41. Li L, Liu W, Zhang H, et al.: A New Method for Programmable RNA Editing Using CRISPR Effector Cas13X.1. Tohoku Journal of Experimental Medicine 260: 51–61, 2023.

42. Calvo SE, Pagliarini DJ and Mootha VK: Upstream open reading frames cause widespread reduction of protein expression and are polymorphic among humans.

43. Brigham BS, Kitzrow JP, Reyes JPC, Musier-Forsyth K and Munro JB: Intrinsic conformational dynamics of the HIV-1 genomic RNA 5’UTR. Proc Natl Acad Sci U S A 116: 10372–10381, 2019.

44. Bou-Nader C, Link KA, Suddala KC, Knutson JR and Zhang J: Structures of complete HIV-1 TAR RNA portray a dynamic platform poised for protein binding and structural remodeling. Nature Communications 16, 2025.

45. Clerzius G, Gélinas JF and Gatignol A: Multiple levels of PKR inhibition during HIV-1 replication. Rev Med Virol 21: 42–53, 2011.

46. Clerzius G, Shaw E, Daher A, et al.: The PKR activator, PACT, becomes a PKR inhibitor during HIV-1 replication. Retrovirology 10, 2013.

47. Bannwarth S and Gatignol A: HIV-1 TAR RNA: The Target of Molecular Interactions Between the Virus and its Host. Curr HIV Res 3: 61–71, 2005.

48. Jean MJ, Zhou D, Fiches G, et al.: Curaxin CBL0137 has the potential to reverse HIV-1 latency. J Med Virol 91: 1571–1576, 2019.

49. Sánchez-Luque FJ, Reyes-Darias JA, Puerta-Fernández E and Berzal-Herranz A: Inhibition of HIV-1 replication and dimerization interference by dual inhibitory RNAs. Molecules 15: 4757–4772, 2010.

50. Ka WH, Jeong YY and You JC: Identification of the HIV-1 packaging RNA sequence (Ψ) as a major determinant for the translation inhibition conferred by the HIV-1 5’ UTR. Biochem Biophys Res Commun 417: 501–507, 2012.

51. Ishimaru D, Plant EP, Sims AC, et al.: RNA dimerization plays a role in ribosomal frameshifting of the SARS coronavirus. Nucleic Acids Res 41: 2594–2608, 2013.

52. Ye L, Gribling-Burrer AS, Bohn P, et al.: Short- and long-range interactions in the HIV-1 5′ UTR regulate genome dimerization and packaging. Nat Struct Mol Biol 29: 306– 319, 2022.

53. Jouvenet N, Lainé S, Vivares LP and Mougel M: Cell biology of retroviral RNA packaging. RNA Biol 8, 2011.

54. Chung CH, Allen AG, Atkins AJ, et al.: Safe CRISPR-Cas9 Inhibition of HIV-1 with High Specificity and Broad-Spectrum Activity by Targeting LTR NF-κB Binding Sites. Mol Ther Nucleic Acids 21: 965–982, 2020.

55. Bobbin ML, Burnett JC and Rossi JJ: RNA interference approaches for treatment of HIV-1 infection. Genome Med 7, 2015.

56. Shortridge MD, Wille PT, Jones AN, et al.: An ultra-high affinity ligand of HIV-1 TAR reveals the RNA structure recognized by P-TEFb. Nucleic Acids Res 47: 1523–1531, 2019.

57. Gatignol A, Buckler-White A, Berkhout B and Jeang K-T: Characterization of a Human TAR RNA-Binding Protein That Activates the HIV-1 LTR. Science (1979) 251: 1597– 1600, 1991.

58. Fujii R, Okamoto M, Aratani S, et al.: A Role of RNA Helicase A in cis-Acting Transactivation Response Element-mediated Transcriptional Regulation of Human Immunodeficiency Virus Type 1. Journal of Biological Chemistry 276: 5445–5451, 2001.

59. Yamaguchi Y, Inukai N, Narita T, Wada T and Handa H: Evidence that Negative Elongation Factor Represses Transcription Elongation through Binding to a DRB Sensitivity-Inducing Factor/RNA Polymerase II Complex and RNA. Mol Cell Biol 22: 2918–2927, 2002.

60. Agbottah ET, Traviss C, McArdle J, Karki S, St. Laurent GC and Kumar A: Nuclear factor 90(NF90) targeted to TAR RNA inhibits transcriptional activation of HIV-1. Retrovirology 4, 2007.

61. Svitkin Y V, Pause,’t And A and Sonenbergl N: La Autoantigen Alleviates Translational Repression by the 5’ Leader Sequence of the Human Immunodeficiency Virus Type 1 mRNA., 1994.

62. Bou-Nader C, Gordon JM, Henderson FE and Zhang J: The search for a PKR code-differential regulation of protein kinase R activity by diverse RNA and protein regulators.

63. Balachandran S, Roberts PC, Brown LE, Truong H, Pattnaik AK, Archer DR and Barber GN: Essential Role for the dsRNA-Dependent Protein Kinase PKR in Innate Immunity to Viral Infection., 2000.

64. Zhu PJ, Huang W, Kalikulov D, et al.: Suppression of PKR promotes network excitability and enhanced cognition by interferon-γ-mediated disinhibition. Cell 147: 1384–1396, 2011.

65. McKenna SA, Kim I, Liu CW and Puglisi JD: Uncoupling of RNA Binding and PKR Kinase Activation by Viral Inhibitor RNAs. J Mol Biol 358: 1270–1285, 2006.

66. Spanggord RJ, Vuyisich M and Beal PA: Identification of Binding Sites for Both dsRBMs of PKR on Kinase-Activating and Kinase-Inhibiting RNA Ligands. Biochemistry 41: 4511–4520, 2002.

67. Hu W, Kaminski R, Yang F, et al.: RNA-directed gene editing specifically eradicates latent and prevents new HIV-1 infection. Proc Natl Acad Sci U S A 111: 11461–11466, 2014.

68. Kumar S, Hsiao YW, Wong VHY, et al.: Characterization of RNA editing and gene therapy with a compact CRISPR-Cas13 in the retina. Proc Natl Acad Sci U S A 121, 2024.

